# The role of the orbitofrontal cortex in creating cognitive maps

**DOI:** 10.1101/2022.01.25.477716

**Authors:** Kauê Machado Costa, Robert Scholz, Kevin Lloyd, Perla Moreno-Castilla, Matthew P. H. Gardner, Peter Dayan, Geoffrey Schoenbaum

## Abstract

We use internal models of the external world to guide behavior, but little is known about how these cognitive maps are *created*. The orbitofrontal cortex (OFC) is typically thought to access these maps to support model-based decision-making, but it has recently been proposed that its critical contribution may be instead to integrate information into existing and new models. We tested between these alternatives using an outcome-specific devaluation task and a high-potency chemogenetic approach. We found that selectively inactivating OFC principal neurons when rats learned distinct cue-outcome associations, but prior to outcome devaluation, disrupted subsequent model-based inference, confirming that the OFC is critical for creating new cognitive maps. However, OFC inactivation surprisingly led to generalized devaluation. Using a novel reinforcement learning framework, we demonstrate that this effect is best explained not by a switch to a model-free system, as would be traditionally assumed, but rather by a circumscribed deficit in defining credit assignment precision during model construction. We conclude that the critical contribution of the OFC to learning is regulating the specificity of associations that comprise cognitive maps.

**One Sentence Summary:** OFC inactivation impairs learning of new specific cue-outcome associations without disrupting model-based learning in general.

## Introduction

Animals behave in ways that suggest that the brain can build, store, and use internal representations that account for the predictive relationships between elements in the external world. Also called associative models or cognitive maps^1^, these mental constructs are thought to be especially important for adaptive behavior under new or changed conditions^2,3^. The inability to use such models properly is thought to be a key feature of mental illnesses such as schizophrenia^4^, substance use disorder^5,6^, and obsessive compulsive disorder^7^. However, despite their importance, we are only beginning to understand the informational structure of cognitive maps and how the brain creates, stores, and uses them.

In this regard, the orbitofrontal cortex (OFC) has been extensively implicated in model-based behaviors ^8–11^. However, its exact contributions to defining or using the cognitive maps that support these behaviors are still controversial. The currently prevailing view is that the OFC accesses information stored elsewhere to represent the current task space at the time a decision is made ^12–15^. While broadly consistent with the literature, this view is most strongly supported by devaluation experiments in which pairing a given outcome with illness (or satiety) leads to reduced conditioned responding to a cue predicting that outcome. This effect has been shown repeatedly and across species to require the OFC at the time of the probe test ^16–19^, a result generally interpreted as showing a role for OFC in using the associative map formed earlier in training. Compromising the OFC disrupts this usage, resulting in supposedly “model-free” or habit-like behavior. By this account, the OFC offers a form of specialized working memory required for mental simulation.

However, recent studies suggest that the OFC might instead serve as the cognitive “cartographer”, playing a critical role not merely in using maps drawn by other regions but rather in creating and modifying the maps on which other regions rely ^20^. According to this view, OFC manipulations in devaluation probes affect behavior not because OFC is required for mental simulation but rather because the test requires changes to, or recombinations of, existing cognitive maps.

A logical, but untested, corollary of this alternative proposal is that the OFC should also be necessary during initial conditioning in the reinforcement devaluation task, when a major part of the cognitive map used in the later probe is *created.* On the other hand, if the classic view is correct – that, at the time of decision-making, the OFC just uses maps made and maintained elsewhere – then this region should not be necessary during the conditioning phase. As one cannot read what is not yet written, this prediction allows for an acid test to differentiate whether the OFC is a reader or a cartographer of cognitive maps. Here, we performed this test using a within-subject outcome-specific devaluation task and high-potency chemogenetics to inactivate OFC transiently when maps were first being formed.

## Results

Food restricted rats, transfected with either hM4d (inhibitory DREADD receptor, n=15) or only mCherry (control; n=13) in the OFC (Figure S1), underwent conditioning in which two different auditory cues (A and B) predicted the delivery of either banana- or bacon-flavored pellets (Figure 1A). Before each session, rats were injected with JHU37160 dihydrochloride (JH60; i.p. 0.2 mg/kg), a high-potency DREADD agonist ^21^, to inactivate OFC principal neurons in hM4d-transfected rats both transiently and selectively ^22^. The use of this new generation compound avoids several confounds associated with other DREADD agonists ^21,23^.

**Figure 1.**
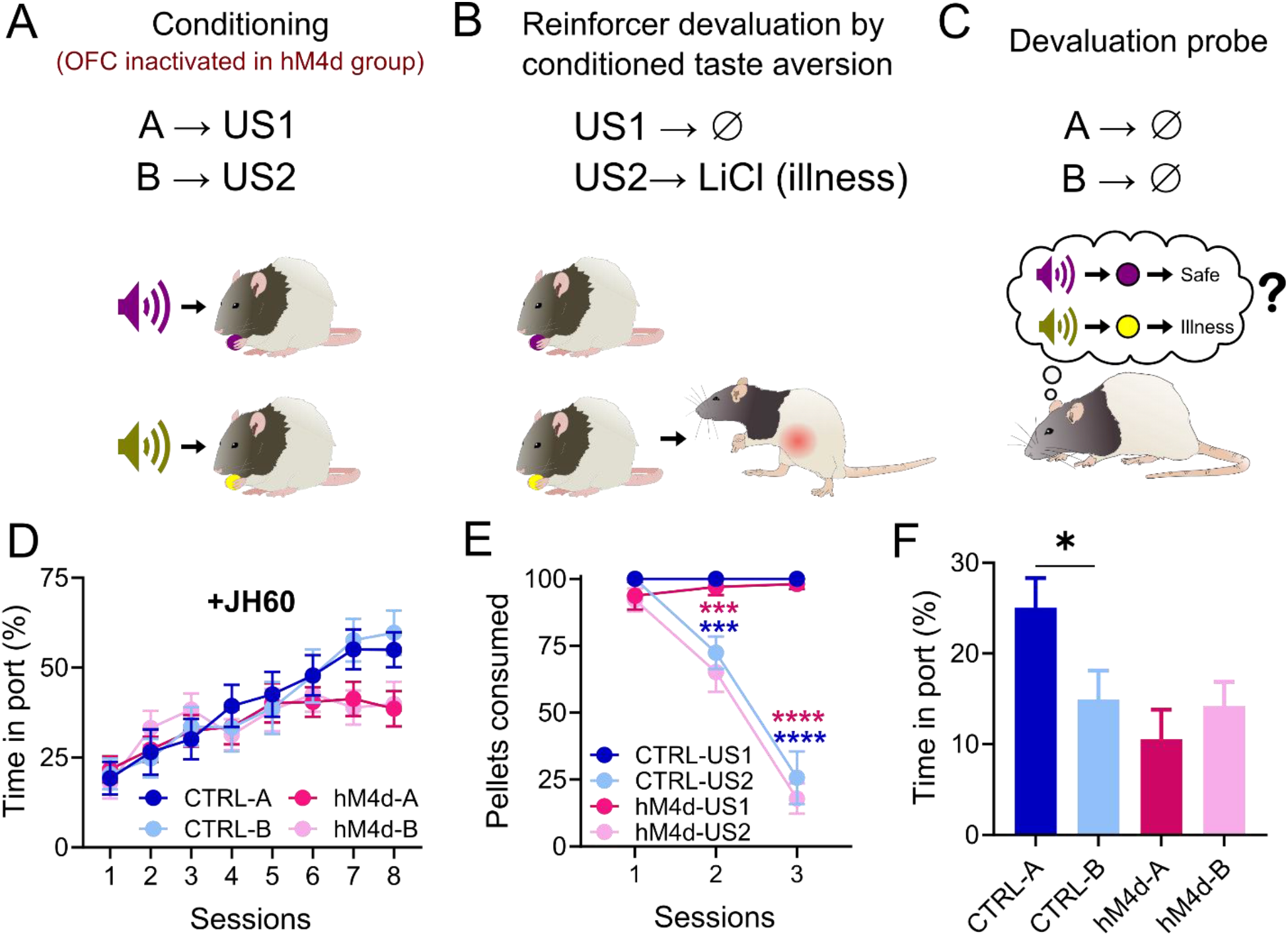
Chemogenetic inactivation of OFC during conditioning abolishes subsequent sensory-specific cue devaluation. **(A-C)**: Schematic of the behavioral procedures. **(A):** Rats were conditioned to two cues, A and B, which lead to different rewards. The OFC was inactivated in the hM4d group. **(B):** Later, one of the rewards was paired with LiCl injections. **(C):** Finally, rats were re-exposed to the conditioned cues, testing if a model-based association had been established between them and the rewards. Food cup responding during conditioning. There was no isolated or interaction effect of cue identity (*P*>0.05), nor an effect of group (*P*>0.05), and rats of both groups increased responding as sessions progressed (*P*<0.0001****). However, there was a significant interaction between group and session progression (*P*<0.0001****), visible in the last two sessions. **(E):** Pellet consumption during CTA. Rats from both groups consumed nearly all pellets in the first CTA session, and consumed less of the devalued pellet type as sessions progressed (*P*<0.0001****). **(F):** Food cup responding during probe. There was a significant effect of group (*P*=0.047*), and the interaction of the group with the cues (*P*=0.009**), as control rats responded more to A than to B, while hM4d rats responded equally to both cues. Asterisks in graphs indicate post-hoc multiple comparison test results. See Table S1 for detailed statistics. Data are represented as mean ± SEM. \**P*<0.05; \*\*\**P*<0.001; \*\*\*\**P*<0.0001.

Despite inactivation, rats in both groups progressively increased responding to the food cup during presentation of either cue (Figure 1D). Initial acquisition rates were similar, although rats in the hM4d group responded slightly less during the last two sessions of conditioning, in agreement with recent work showing that transient OFC inactivation can reduce asymptotic conditioned responding in some settings ^24^.

After conditioning, rats were subjected to conditioned taste aversion (CTA) training, in which one of the rewards (the one associated with B), was paired with LiCl injections, inducing nausea (Figure 1B). Rats initially preferred both rewards equally, but quickly and selectively reduced consumption of the pellet type paired with LiCl (Figure 1E).

Finally, after CTA training, rats were given a probe test, in which the cues were presented as during conditioning but without reward (Figure 1C). As expected, control rats responded more to cue A (paired with the non-devalued pellet) than to cue B (paired with the devalued pellet), indicating they had learned the specific cue-reward and reward-illness associations and were able to integrate them in the probe test to infer that B might lead to devalued reward (Figure 1F). By contrast, rats in the hM4d group responded equally to both cues (Figure 1F). This result is inconsistent with the hypothesis that OFC’s main function is to access mental maps stored elsewhere to support model-based behaviors at the time a decision is made, and instead supports the alternative hypothesis that OFC plays a critical role in drawing those maps during initial learning ^20^.

That said, while this result supports this alternative hypothesis, rats in the hM4d group did not simply lack the devaluation effect, as would be expected if there was no model, but rather they appeared to generalize the devaluation effect across cues (Figure 1F). This was evident even if responses during the probe were normalized to the end of conditioning, indicating that the effect was not related to modest reduction in asymptotic conditioned responding (Figure 2A). That the two effects were orthogonal to each other is further supported by the lack of correlation between responding at the end of conditioning and the effect of devaluation (Figure 2B). Nor was the apparent generalization due to differences in CTA retention as preference tests revealed that CTA effects were similar in the two groups after the probe test (Figure 2C).

**Figure 2.**
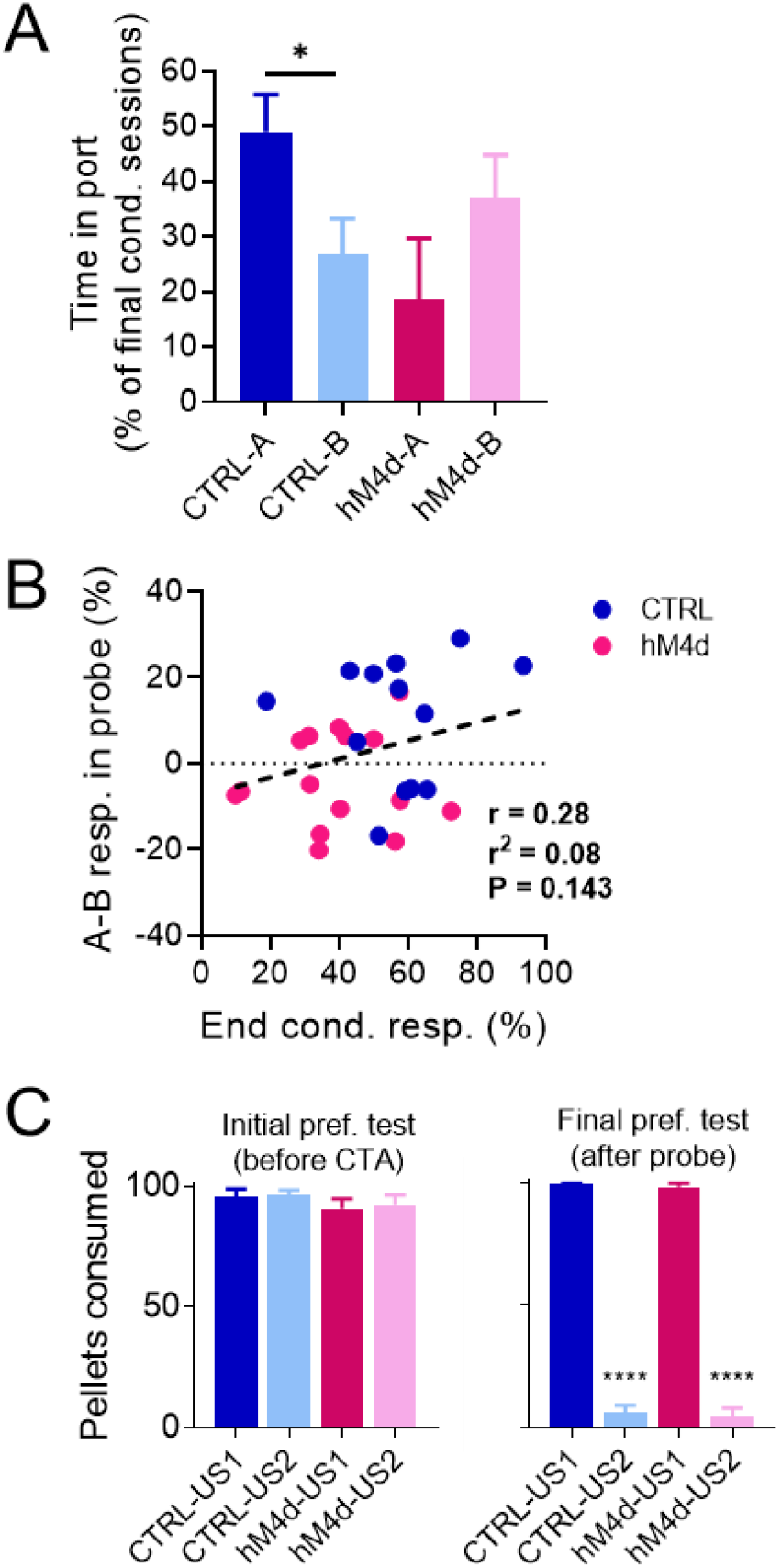
Generalization of devaluation due to OFC inactivation is not dependent on effects on conditioned responding or CTA learning. **(A):** Food port responding in the final probe session but normalized to the last two sessions of conditioning. There was a significant interaction effect of the group with the cues *(P* = 0.002**), as well as only a significant difference between A and B in the control group in the post-hoc test. **(B):** Differential responding to valued and devalued cues (responding to A – mean responding to B) was not correlated to the conditioned responding at the end of initial learning (average of % time in port for both cues in the last two sessions of conditioning). **(C):** Consumption of pellets during preference tests for CTRL (blue and light blue) and hM4d (red and pink) rats. Rats from both groups consumed all pellets similarly during the first preference test (2-way ANOVA; ND x D: *F*_1,26_ = 0.12, *P* = 0.7318; CTRL vs hM4d: *F*_1,26_ = 1.235, *P* = 0.2766; interaction: *F*_1,26_ = 0.0171, *P* = 0.8969) and both groups equally consumed significantly less of the devalued pellet type (the one previously associated with cue B and paired with LiCl during CTA) in the second preference test (2-way ANOVA; ND x D: *F*_1,26_ = 1364, *P* < 0.0001****; CTRL vs hM4d: *F*_1,26_ = 0.3519, *P* = 0.5582; interaction: *F*_1,26_ = 0.0005, *P* = 0.9825). Asterisks in the graphs indicate results of post-hoc multiple comparison tests. Data are represented as mean ± SEM. \*\*\*\**P*<0.0001.

Generalization of devaluation also could not be accounted for by effects of OFC inactivation on perception or memory. To show this, we tested a subset of these rats in an object recognition task ^25^. OFC was inactivated prior to the sample phase of the task, while the rats first explored two identical objects (Figure 3A). Over the next 2 days, the rats were brought back to the same arena for two recognition tests in which novel objects were substituted for the objects introduced in the sample phase (Figure 3B-C). If OFC inactivation in these rats induced perceptual confusion, accelerated forgetting, or context-dependent learning, then inactivation in the sample phase of this task should have disrupted object discrimination in the first but not the second recognition test, yet we found no such effect (Figure 3D-I).

**Figure 3.**
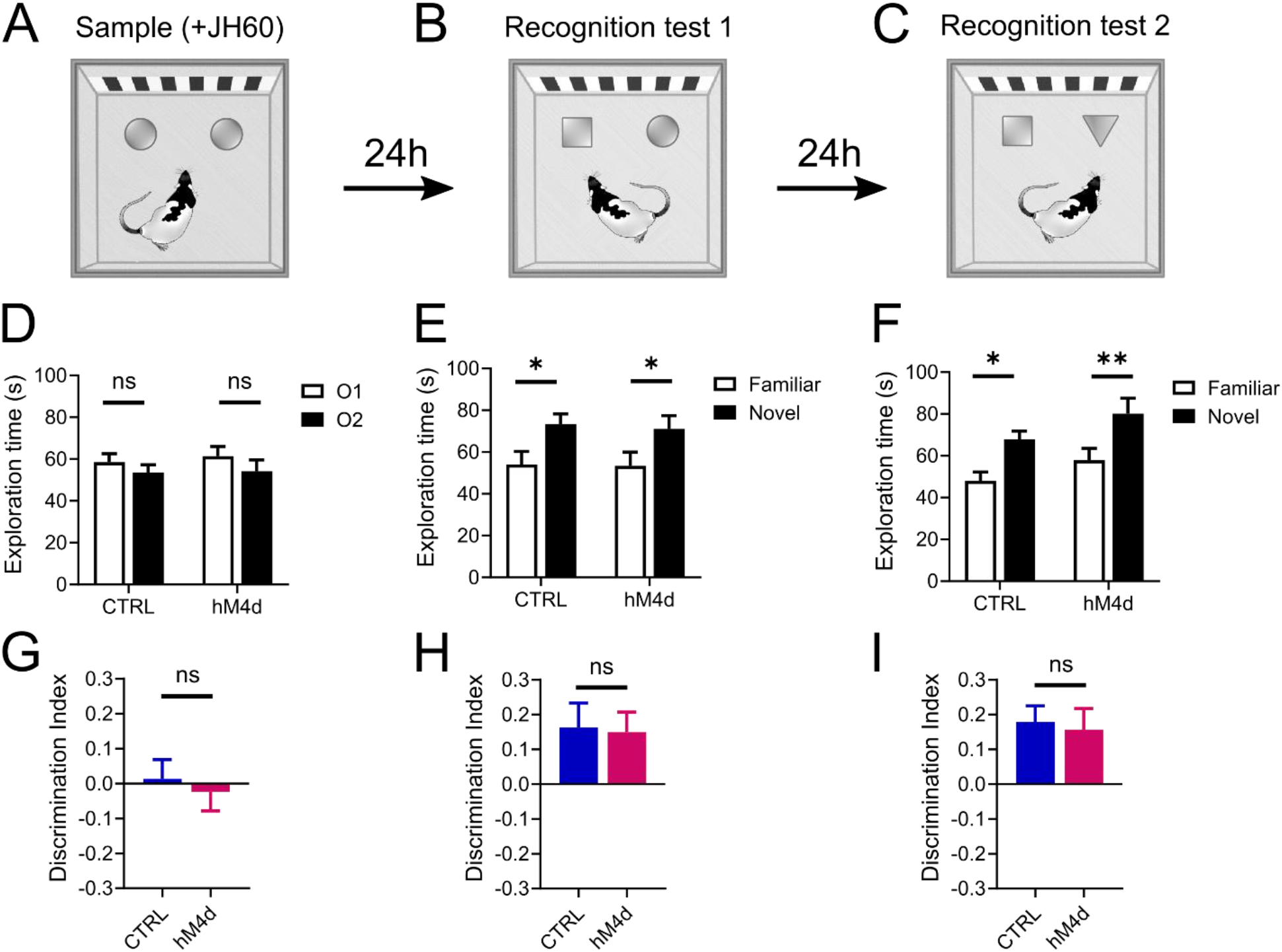
OFC inactivation does not affect object recognition. **(A):** Sample phase, where rats explored two identical objects and received JH60 injections. **(B):** First recognition test, where one familiar object was replaced by a novel one. **(C):**Second recognition test, where the previous familiar object was substituted by yet another novel object. **(D and G):** Rats in both groups explored the two objects for the same amount of time during sample (2-way ANOVA; O1 x O2: *F*_1,17_ = 1.833, *P* = 0.193; CTRL vs hM4d: *F*_1,17_ = 0.14, *P* = 0.712; interaction: *F*_1,26_ = 0.059, *P* = 0.809)(D) which was evident in the discrimination index (unpaired t-test, *P* = 0.634)(G), demonstrating that OFC inactivation does not affect exploratory behavior in this task. **(E and H)**: Rats from both groups showed equally robust object recognition learning, evident in the increased exploration of the novel object (Familiar x Novel: *F*_1,17_ = 13.53, *P* = 0.002**; CTRL vs hM4d: *F*_1,17_ = 0.045, *P* = 0.835; interaction: *F*_1,26_ = 0.025, *P* = 0.876) (E) and an increase in the discrimination index, which was identical between groups (*P* = 0.882) (H), indicating that OFC inactivation in sample did not affect recognition learning or memory retention, nor did it induce some form of context-dependent learning. **(F and I):** Again, rats in both the control and hM4d groups showed a similar level of preference for the novel object (Familiar x Novel: *F*_1,17_ = 18.13, *P* = 0.0005***; CTRL vs hM4d: *F*_1,17_ = 3.085, *P* = 0.097; interaction: *F*_1,26_ = 0.053, *P* = 0.82) (F), as confirmed in the discrimination index (*P* = 0.775)(I), confirming that learning under the effects of JH60 injections was similar to when no drug was injected. Asterisks in E and F indicate results of post-hoc multiple comparison tests. Data are represented as mean ± SEM. \**P*<0.05, \*\**P*<0.01.

The generalization of devaluation in the OFC inactivated group was unexpected and intriguing, since model-based learning is traditionally treated as an all-or-none phenomenon. A complete failure of model-based control would leave only devaluation-insensitive, model-free behavior intact, resulting in high responding to both cues. It has been proposed that associative learning may operate as a dynamic mixture of model-based and model-free learning ^3^, and that the OFC may mediate this process ^26^. Therefore, we considered whether our results could be explained by a change in the balance between model-based to model-free learning under OFC inactivation. This explanation has some intrinsic disadvantages, as it requires at least two parallel learning systems and a third process to integrate their outputs, i.e., it is complex, with many free parameters. We found that it was possible to reproduce our results with this approach provided we also added a forgetting parameter (Figures S3). However, the resultant fits were hard to reconcile with the general understanding of OFC function, as they did not produce a decrease in model-based learning with OFC inactivation, but rather an increase in model-free learning rates (Figure S3C). This suggests a form of structural over-fitting, consistent with the observation that the fitted parameters could not be reliably recovered from simulated data (Figure S3D). Thus, a complete or partial shift from model-based to model-free control seemed not to offer a good explanation for the experimental results.

A more promising way to account for the results is to consider the possibility that the subjects are still building, and then using, a cognitive map, but that the map is different – perhaps less precise – without the contribution of OFC. This idea would be consistent with recent arguments against pure model-free processing ^27,28^, evidence that the OFC is particularly important for sculpting representations of various aspects of tasks^13,29,30^, and findings in OFC-lesioned macaques of impaired credit assignment ^31^. Translating this idea to the current task, we hypothesized that the OFC might be particularly important for segregating and separately updating each unique cue-outcome pair, which were of uncertain importance in initial conditioning.

We tested this proposal by fitting our data with a novel model-based reinforcement learning algorithm trained on the same sequence of trials as in the task ^3,32^ (Figure 4). The effect of OFC inactivation on learning during initial conditioning was captured by introducing an “imprecision” parameter (*χ*) that defined how credit assignment spread – i.e., whether updates were selective for each cue-outcome pair during the conditioning phase of the task (Figure 4A). Thus, receiving a banana-flavored pellet after cue A updates the association between the alternative cue B and the banana-flavored pellet by an amount proportional to *χ*. Only if *χ* = 0, would the update be confined exclusively to cue A. A model with a high *χ* value would therefore be able to learn that auditory cues predict sucrose pellets, but would have trouble differentiating which pellet flavor (e.g., banana) is associated with which cue (A or B). Substantial confusion during conditioning (high *χ*) would cause the loss of value imposed by the following CTA training (Figure 4B) to be at least partially generalized to both cues A and B, due to the imprecision of specific state predictions and subsequent inference (Figure 4C), noting that the rats remained well aware of the separate values of the pellet types after the probe test (Figure 2C, right).

**Figure 4.**
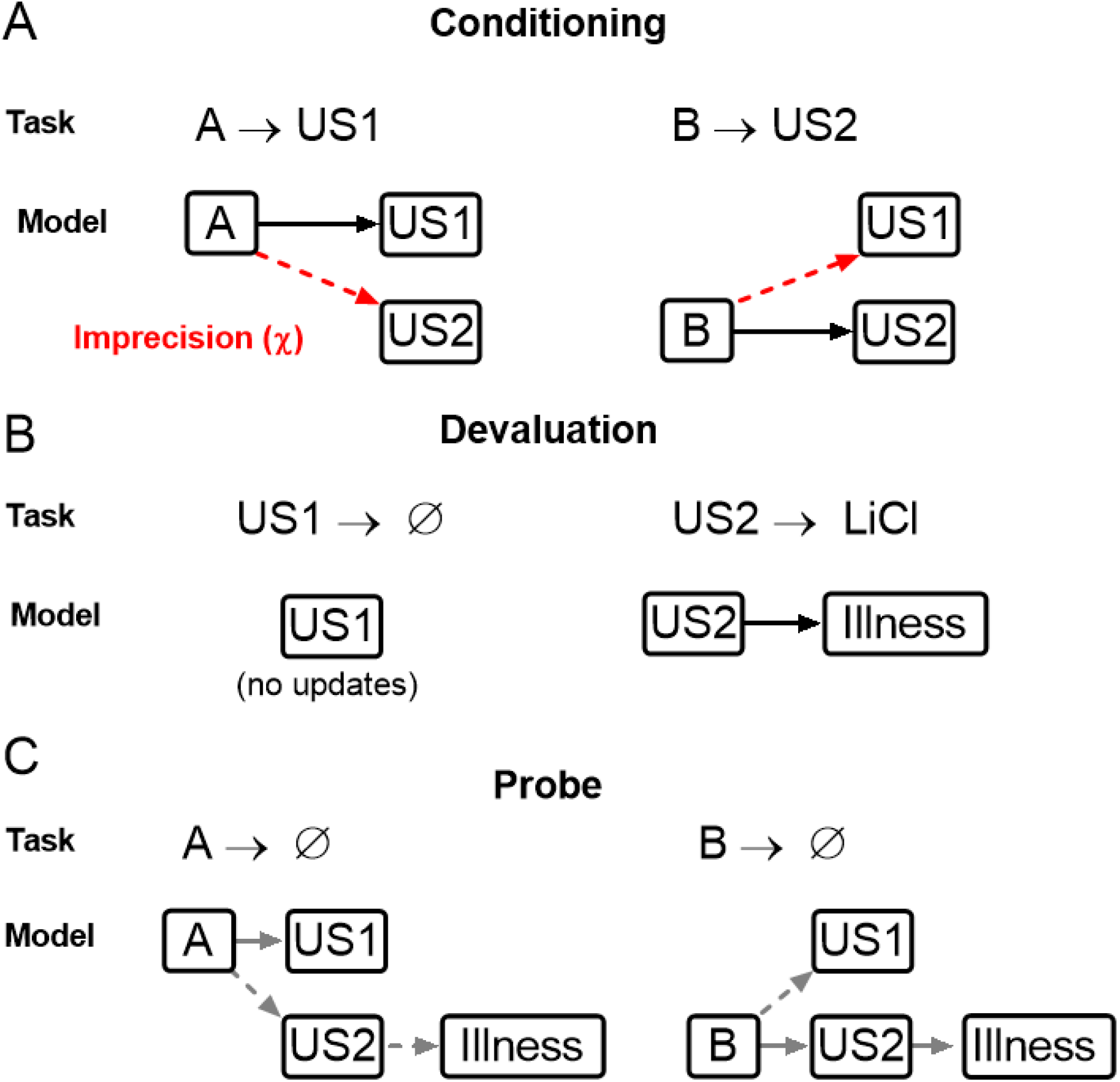
A model-based reinforcement learning algorithm that simulates imprecise state identity credit assignment. **(A):** During initial conditioning, the value and state transition matrices are updated according to the task contingencies (A-R1, B-R2; solid black arrows), except for a parallel updating of the opposite association (A-R2, B-R1), which occurs proportionately to the imprecision term *χ* (dashed red arrows). **(B):** During the CTA devaluation procedure, updating obeys task contingencies, with no value or state prediction updates to the R1 state, but with a learning that R2 predicts a devalued illness state. **(C)** During the probe, new learning follows the task states, and the value of cue states is adjusted according to the inferred state predictions (grey arrows), including generalized inferences driven by the imprecision term during initial acquisition (dashed grey arrows).

We found that this “imprecision” model fit our behavioral results well (Figure 5), reproducing the normal behavior in the control group and all effects of OFC inactivation, including both the apparent generalization of devaluation in the probe test (Figure 5B-C) as well as the lower asymptotic performance in conditioning (Figure 5E-F). Critical parameters in the model, particularly *χ,* were recoverable from simulated data (Figure S2A) ^33,34^. Model fits to data from control and hM4d groups differed in their imprecision term *χ,* which was significantly higher in hM4d models (Figure 5B and Table S2). Furthermore, *χ* was highly correlated with the difference in responding to the valued (A) versus devalued (B) cues during probe (Figure 5C), even though this parameter only affected learning during conditioning (Figure 4A). Notably, this effect was not due to an effect of *χ* on the strength of conditioning, as these were uncorrelated (Figure 5D).

**Figure 5.**
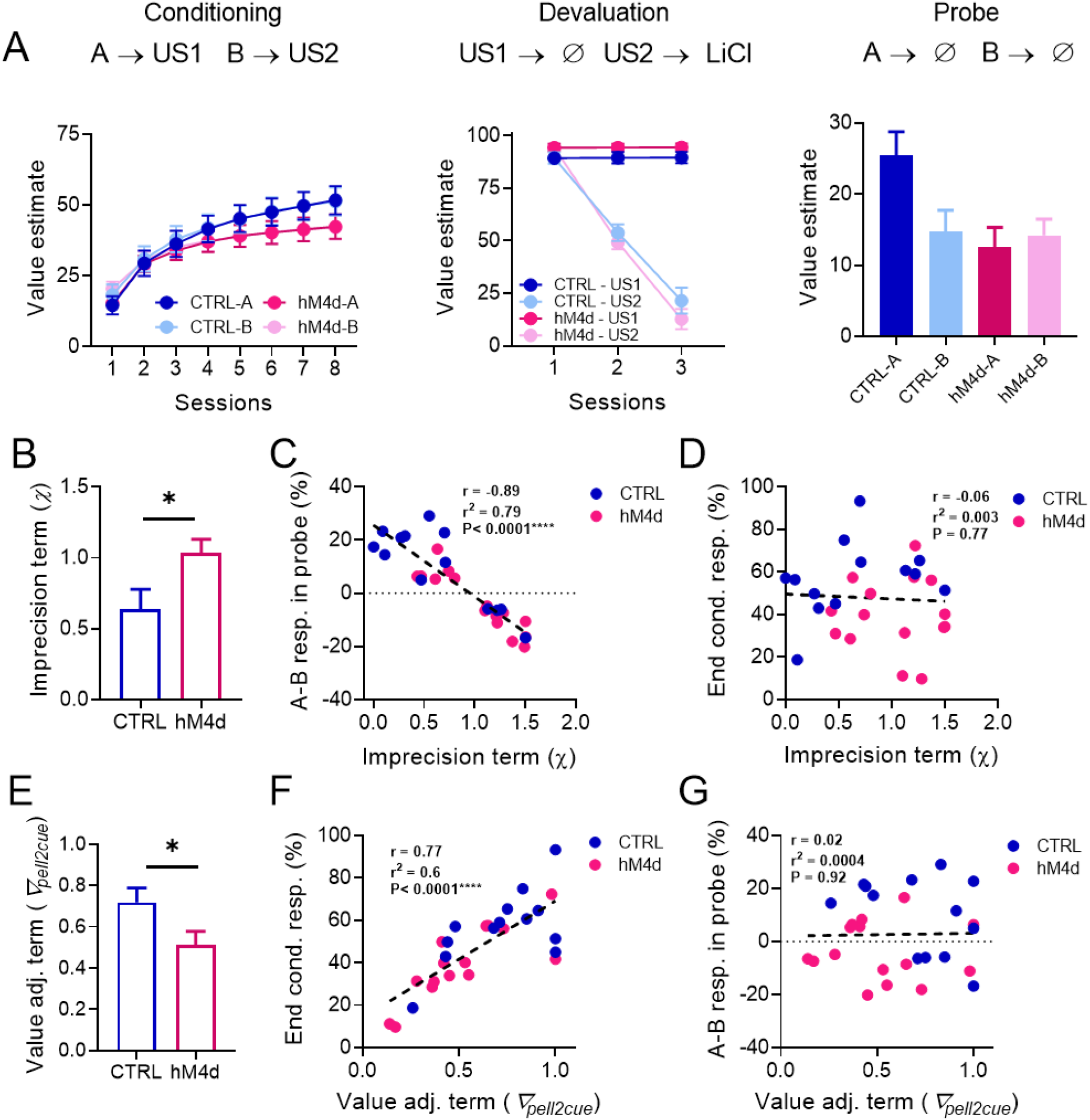
OFC inactivation effects on reinforcer devaluation are explained by a deficit in differentiating specific cue-outcome associations. **(A):** Model fit results for our model-based reinforcement learning model with potential outcome identity confusion. **(B):** The imprecision term *χ* was significantly higher in models fitted to hM4d behavioral data in relation to controls (*P*=0.027*). **(C):** *χ* was negatively correlated with the differential responding to cues in the probe session. **(D):** *χ* was not correlated with the average responding to cues at the end of conditioning. **(E)**: The value adjustment term ∇_pell2cue_ was significantly lower in hM4d models (*P* = 0.04*). **(F):** ∇_pell2cue_ was positively correlated with average cue responding at the end of conditioning. **(G)**: ∇_pell2cue_ was uncorrelated with differential responding to cues in the probe session. See Table S2 and Figure S2 for detailed parameter comparisons. Data are represented as mean ± SEM. \**P*<0.05.

Our model also recapitulated other aspects of the results, specifically by having a value adjustment parameter (*∇*_pell2cue_) that captured the asymptotic performance during conditioning. The value of this parameter differed between fits for control and hM4d subjects (Figure 5E), accounting for the reduced responding of hM4d rats at the end of conditioning (Figure 1D, 2B and 5F). Importantly, *∇*_pell2cue_ did not correlate with the difference in cue responses during the probe (Figure 5G). These results confirm that the effects of OFC inactivation during model creation on subsequent model-based decision making are not related to the concurrent effects on asymptotic value estimation. The latter may be related to the known role of OFC in representing and updating outcome value ^14,35^.

## Discussion

Our study demonstrates first that OFC is necessary for the construction of a normal cognitive map and second that the OFC appears to play a circumscribed role in this construction process. In our task, the map-making apparently did not cease when OFC was inactivated, but the created map was degraded and less specific about which cues led to which outcomes. This was modeled as a lack of precision in credit assignment, but a failure to create appropriately “granular” ^36,37^ internal representations of these external events would produce the same result and seems more likely than a direct control of OFC over error signal assignment.

As an intuitive example of the utility of setting this granularity properly, an older child may learn that McDonald’s™ serves Happy Meals™ while Burger King™ serves King Jr™ meals, each with different toys, while a younger sibling may only recall that fast food restaurants serve kids’ meals. Both cognitive maps lead to food, but only one will help you collect all the Disney™ dragons! Whether to keep or discard the information related to which restaurant serves which kids’ meal with which toy is a question of how to segregate the states during learning^36,37^; it is this process that we propose OFC controls or contributes to during cognitive mapping ^20^. This example also illustrates the fact that the generalization afforded by discarding information is not automatically incorrect – it should respond to the exigencies of the circumstance.

We would speculate that OFC’s particular contribution to this process is in determining whether to maintain separation between states that have uncertain or perhaps only potential biological significance, when other parts of the circuit might collapse them. Maps formed with too little separation due to hypofunction in OFC would tend to underrepresent potential or hidden associations and meaning and be unable to link to and infer relationships to other maps, as we have seen here. This is also evident in substance use disorder, neurodegenerative diseases, and advanced aging, in which OFC function is compromised^5,6,38–41^, and in children and adolescents, which have immature frontal cortices^42,43^. Conversely, maps formed with too much separation due to an over-exhuberant OFC would tend to instill hidden meaning where it does not exist; notably such an effect is arguably evident in obsessive compulsive disorder and paranoid psychosis, which involve hyperfunction in the OFC and related areas^7,40,44–49^.

The proposal that the OFC plays a critical role in defining the states that form the basis of cognitive maps is congruent with much existing data. This includes classic findings based on manipulations in the probe phase of reinforcer devaluation experiments^16–19,50^, since the probe phase confounds the integration of established maps with their first time use. That is, the function proposed here would be invoked in the probe test in devaluation by the need to recognize the common reward state in the maps created during the conditioning and devaluation phases. Similar conclusions apply to other cardinal studies that have implicated the OFC in model-based behaviors, since these also normally involve integrating or changing task maps ^51–53^. This more limited role for OFC also explains better why this area is necessary in many other behavioral settings where normal behavior depends upon recognizing states that are somewhat ambiguously defined with regard to biological value, including for instance the differential outcomes effect^54^, latent inhibition ^22^, and reversal learning^55–57^, and why OFC seems to grow less important in settings like reversal learning or economic choice once maps are well-established^17,20,55,58^ Finally, perhaps the most intriguing implication of our finding that OFC inactivation fails to reveal model-free learning is the possibility that most learning is, to some degree, model-based, but that mental representations or cognitive maps can be formed with different degrees of granularity or specificity, as determined by the circuits that are engaged in the learning process, including the OFC and other prefrontal areas. In the absence of experimental interventions, illness, or lesions, it could be that the main determinant of the resolution of a cognitive map would be task requirements and learning context. This would mean that perhaps there is a unified learning process that can be more or less complex depending on the contribution of specific circuits or environmental demands.

## Materials and Methods

### Experimental Model and Subject Details

Experiments were performed on 32 male Long-Evans rats (n=16 for each group, >3 months of age at the start of the experiment, Charles River Laboratories) housed on a 12 hr light/dark cycle at 25 °C. Rats were food restricted to ~85% of their original weight for the duration of the experiments and were tested at the NIDA-IRP in accordance with NIH guidelines determined by the Animal Care and Use Committee. All rats had *ad libitum* access to water during the experiment and were fed 16-20 g of food per day, including rat chow and pellets consumed during the behavioral task. Behavior was performed during the light phase of the light/dark schedule. The number of rats used was determined based on previous publications from the lab using Pavlovian conditioning tasks. Prior to surgery, rats were handled every other day for 5-10 minutes for one week. Handling procedures included the performance of mock i.p. injections (rats were scruffed and the experimenter gently poked their abdomen with his finger or the end of a syringe with no needle attached) to prepare the subjects for future real injections. These rats were also used in another study ^22^. One rat in each group was excluded due to incorrect anatomical placement, and two rats were excluded from the control group due to a hardware malfunction during one of the behavioral sessions, leading to n=13 for the control group and n=15 for the hM4d group.

### Surgical procedures

Rats were anesthetized with 1-2% isoflurane and received either AAV8-CaMKIIa-hM4d-mCherry (a Gi-coupled designer receptor exclusively activated by designer drugs (DREADD)) or AAV8-hSyn-mCherry (control), both purchased from Adgene (Cambridge, MA), bilaterally into the OFC (AP −3.0 mm, ML ± 3.2 mm, and DV −4.4 and −4.5 mm from the brain surface) ^22^. A total 0.5 μL was delivered in each site at 0.1 μL/min via an infusion pump.

### Sensory-specific conditioning

Rats were trained and tested at least eight weeks after the surgeries in standard behavioral boxes (12” x 10” x 12,” Coulbourn Instruments, Holliston, MA). Each box was equipped with a food cup, a pellet dispenser and two wall speakers. Head entries into the food cup was measured based on breaks of an infra-red beam.

Rats were conditioned for eight sessions. Prior to each session, each rat received an i.p. injection of JH60 (0.2 mg/kg, dissolved in 0.9% NaCl) and was left in their home cage for at least 15 minutes before the start of the session, to allow for the DREADD agonist to effectively inhibit transfected OFC neurons in the hM4d group ^21,22^.

In every session, rats were exposed to two auditory stimuli, A and B (siren or white noise, counterbalanced across rats); each cue was presented for 10 seconds, immediately followed by the delivery of two bacon- or banana-flavored pellets (TestDiet; counterbalanced pairing). Each pairing was presented eight times per session with an average ITI of 2.5 minutes and the order of presentation was randomized and counterbalanced.

Behavioral responses were quantified as the percentage of time that each rat spent in the food cup during the last 5 seconds of each CS, subtracted by the time they spent in the food cup 5 seconds before CS onset.

### Reward preference tests

Prior to the devaluation procedure, rats were given a preference test comparing consumption of the two pellet-types. Rats were provided 100 pellets of each type, placed in two ceramic bowls for 30 minutes with the location of the bowls reversed every 5 minutes. The remaining pellets were counted after the 30 minute period. This procedure was repeated after the devaluation probe to confirm the permanence of conditioned taste aversion.

### Reinforcer devaluation via conditioned taste aversion with LiCl

For outcome-specific reinforcer devaluation, we paired the reward associated with cue B with LiCl, while the reward associated with cue A was not paired with anything. This devaluation procedure lasted a total of six days. On days 1, 3 and 5, rats were given 30 minutes of access to the devalued pellet, followed immediately by an i.p. injection of 0.3 M LiCl, then returned to their home cages ^17^. On alternate days (2, 4 and 6), rats were given 30 minutes of access to the non-devalued pellet and then returned to their home cages. All preference and consumption tests were performed in clean home cages.

### Devaluation probe

The devaluation probe was performed and analyzed exactly like one of the conditioning sessions, except that no reinforcer was delivered, and the rats did not receive an injection.

### Object recognition task

A subset of 10 rats from each group of the previous experiment was randomly selected for this procedure. One of the control rats was the one excluded due to incorrect anatomical placement, leading to n=9 for the control group and n=10 for the hM4d group for this experiment.

One square arena (60 x 60 cm) made of brown plexiglass with a striped black and white rectangular spatial cue was placed in a dimly (~3 lumens) red-light illuminated room. A video camera was mounted above the arenas, and activity during test sessions was digitized with a high-definition webcam (C920S PRO HD, Logitech, Suzhou, China). The objects to be discriminated were white glass bulbs, transparent glass jars, cylindrical amber glass bottles and trapezoidal white plastic bottles. All objects were glued to heavy metal disks to prevent them from being displaced by the rats, and positioned at the back corners of the arena (10 cm from walls). To avoid olfactory cues, the arena and objects were thoroughly cleaned with 0.1% acetic acid after each trial.

For habituation, the rats were positioned into the open-field arena without any objects for 10 min the day before the start of the experiment. Throughout the experiment, the position of the objects was constant, but the objects used and their relative positions were counterbalanced for every animal. In the sample phase, rats were placed in the arena facing the wall opposite the objects and were allowed to freely explore two identical objects (either two light bulbs or two jars) for 10 min. Prior to the sampling session, each rat received an i.p. injection of JH60 (0.2 mg/kg, dissolved in 0.9% NaCl) and was left in their home cage for at least 20 minutes before the start of the session. This period was given to allow for the DREADD agonist to reach the brain and effectively inhibit transfected OFC neurons in the hM4d group. After 24 h, on memory test 1, rats were allowed to explore freely one copy of the previously presented object (familiar) together with a new one (novel) for 10 min. A second memory test was performed 24 h after the first test. During the second memory test, the object that was introduced in the previous memory test was kept in place (so now it was the familiar object), and the previous familiar object was replaced by a novel object (either amber or white bottles), and rats explored freely for 10 min.

As previously described^25^, exploration was defined as pointing the nose toward to an object at a distance of less than 1 cm and/or touching it with the nose. Turning around or sitting on the objects was not considered as exploratory behavior. A Discrimination Index (DI) was calculated, where DI = difference between exploration of the novel and familiar objects / total object exploration time during each memory test, such as that a DI of 0 indicates equal preference for both objects, a DI of 1 indicates exclusive exploration of the novel object, and a DI of −1 indicates exclusive exploration of the familiar object.This measure was also calculated using only the first 5 min of each test, but results were similar to when the whole test period was used (data not shown). Video recordings were scored automatically using TopScan Suite (Clever Sys, Reston, VA). Exploration times were verified manually by a trained rater blinded to treatment and objects identities using BORIS software (Version 7.9.19, University of Torino, Italy).

### Histological procedures

After completion of the experiment, rats were perfused with chilled phosphate buffer saline (PBS) followed by 4% paraformaldehyde in PBS. The brains were then immersed in 18% sucrose in PBS for at least 24 hours and frozen. The brains were sliced at 40 μm and stained with DAPI (Vectashield-DAPI, Vector Lab, Burlingame, CA). Fluorescent microscopy images of the slides were acquired with a BZ-X800 Keyence microscope. Expression patterns were extracted from the images and then superimposed on anatomical templates ^22^.

### Statistical analyses

Data were analyzed using GraphPad Prism (GraphPad Software, San Diego, CA). Error bars in figures denote the standard error of the mean. Effects of experimental variables on behavioral measures were examined with repeated-measures 2-way and 3-way ANOVAs combined with Sidak’s or Tukey’s post-hoc tests, respectively. Statistical significance threshold for all tests was set at *P*<0.05.

### Reinforcement learning modelling

#### Background

We modelled the five stages of the experiment in chronological order: Conditioning (COND), Preference Test 1 (PRFT1), Devaluation (DEV), Preference Test 2 (PRFT2) and finally Probe testing (PROBE). For COND and PROBE, the *Port Stay Probability (PSP)* upon cue presentation was quantified. In PRFT1, DEV and PRFT2, the *percentage of pellets eaten (PPE)* was quantified. Two pellets of a single type were delivered in each case.

On each trial, an internal value estimate 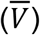 was calculated based on contributions from a model based (MB) system (and, for the alternative hypothesis of a loss of MB learning, in combination with a model free, MF, system). This value estimate was then transformed to the behavioral measurement that was appropriate to the experimental stage. In keeping with standard practice, we described the Pavlovian connection between cue and outcome as being associations; however, in keeping with the temporal evolution of the task, we actual model them as transitions from cue to outcome. MB (and MF) systems were updated using the state transitions that were observed (e.g., A→ValuedOutcome) and the rewards that were received.

The main hypothesis (we call this Ha) that we tested was that the OFC enables precise credit assignment through separation of specific cue-outcome relations (i.e., that sound A predicts banana flavored pallets) and when deactivated, only the general relation (that any auditory cue predicts delivery of food) can be learned. However, we also tested a model (Hb) which could potentially characterize a more conventional view of OFC deactivation, namely that it would suppress MB over MF control. Since Hb mostly nests Ha, we provide an partly integrated discussion.

#### Formal model

*S* = {*s*_1_,...,*s_n_*} is the set of states. Each state is typically associated with the presentation of a cue or an outcome that can be rewarded or devalued, i.e., *S* ~ {*A,B*,ValuedOutcome, DevaluedOutcome}.

In order to be able to characterize MB and MF systems fairly, we considered forms of both that represent the uncertainty in their predictions of rewards and values. However, we adopt a heuristic Bayesian scheme, with observation rates (the equivalent of learning rates) that are parameters (rather than pure conjugate distributional updates).

Following Dearden et al. ^32^, normal-gamma distributions are used to characterize this uncertainty (since, following Daw et al. ^3^, MB and MF systems share the characterization of the values of the final, reward, states, albeit potentially with different parameters, and with only the MB system being subject to the effects of devaluation).

We write this down in terms of the value of state s. The normal-gamma distribution for the value *V_s_* and the *precision* 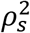 is written as 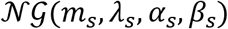. According to this, the *conditional* distribution of *V_s_* given 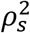 is a normal distribution

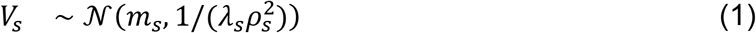

and the precision has an unconditional gamma distribution

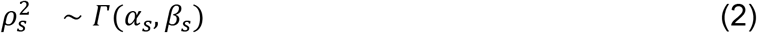

in terms of our problem, we interpret the parameters as follows:

*m_s_* is the mean reward across the previous iterations
*λ_s_* is the number of outcomes seen (this also includes the cases when no reward (*r* = 0) is delivered in this state)
*α_s_* describes the total opportunity for learning about the precision; assuming that we initialize alpha to: 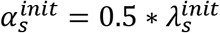, then it holds that at all times *α_s_* = 0.5 * *λ_s_*.
*β_s_* describes the scale of the precision across previous seen rewards

implying that the marginal mean and variance of *V_s_* are:

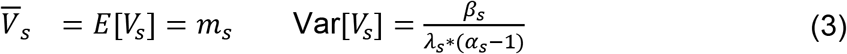

For MB computations, we also need an internal model of the state graph. We use *T* to describe the distribution of transition probabilities from all to all states. Programmatically, *T* can be described by a matrix where each row contains *ϕ*’s that are parameters for the multinomial distribution that characterizes the transition probabilities from a “source” state s to any of the other states (including the source state itself):

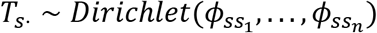

This will only be interpretable for non-terminal “source” states *s*, as the trial ends afterwards and no information about consecutive states can be collected. The terminal states are thus absorbing. The sum of probabilities for a fixed source state to all possible target states is 1 (see model based value calculation).

#### Initialization

We initialize all *ϕ*’s in *T* to 10. This implies a moderately strong prior that the transition probabilities are uniform across all states:

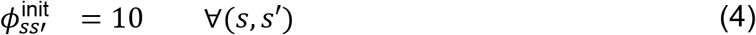

We initialize the distribution describing the reward distribution parameters to:

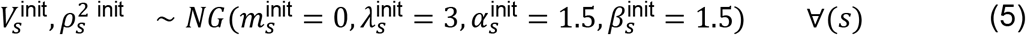

The rationale for these values is that 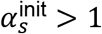 to ensure *V_s_* has a finite marginal variance. The value of 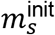 was chosen to be 0 as animals start out with no value expectation. 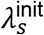 was set to 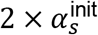, as this ratio is also maintained by the updates. 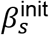 was set to 1.5 in order to set the starting marginal variance to 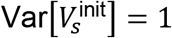. However, we confirmed that our results are stable to quite a wide range of initialization values, provided that the variance is well-defined (*α_s_* > 1).

During the conditioning stage, *r*_ValuedOutcome_ = *r*_DevaluedOuteome_ = 2 (for the number of pellets provided). The reward of the DevaluedOutcome changes during the devaluation period to NegRew < 0, which is a parameter that captures the strength of the devaluation effect for each animal.

#### Model updates and value calculation

The normal-gamma distribution characterizing the value *V_s_* of a terminal state updates according to each observation. In general, given an observation 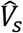, writing 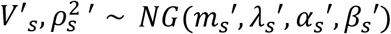 for the updated distribution at s, we update the parameters as:

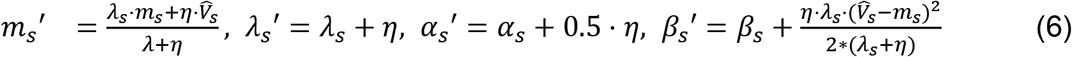

where *η* is called an observation rate and stands in for the number of subjective observations associated with each experience – it need only be positive and is not constrained to be less than 1.

For the MB system, writing 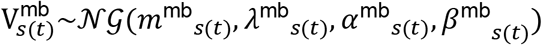, for a terminal state, the update happens using 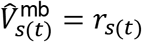 and observation rate *η* = *η*^mb^.

For the transition matrix, if the state *s*(*t*) is a non-terminal state that is followed by state *s*(*t* + 1), the parameters of the transition probability distribution *T*_*s*(*t*)_. are updated using a notional transition observation rate *η*^t^ as:

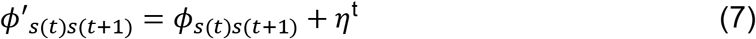

The MB system combines its knowledge of transitions and immediate rewards by applying the Bellman equation, which, in this case is very straightforward, since there are only two steps. Ignoring any posterior correlation between *T* and *μ, σ,* this implies that:

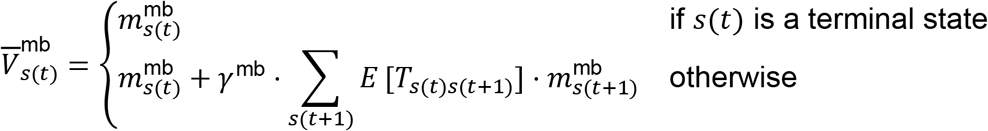

The expected value for the next state is discounted by *γ*^mb^, which normally is close to 1. The expected value for the transition probability from state *s*(*t*) to state *s*(*t* + 1) can be calculated using: – *E*[*T*_*s*(*t*)*s*(*t*+1)_] = *ϕ*_*s*(*t*)*s*(*t*+1)_/∑_ω_*ϕ*_*s*(*t*)*ω*_.

The approximate variance can be calculated from the Bellman equation (again ignoring correlations).

#### Transformation of Estimated Values to Behavioral Measures

Having generated a prediction 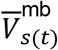 from the MB system, it is necessary to convert it into the different experimental measures used in the various stages of the experimental paradigm. To do this, the combined value is normalized by the standard scalar reward received (2, for the number of pellets), and thresholded at 0 in order to avoid negative percentages when calculating the behavioral measures:

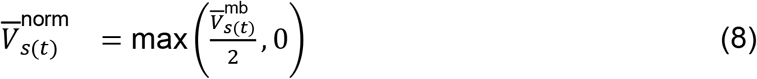

This normalized value can then be transformed to the respective behavioral measures for each stage, each given as percentages in the range [0,100]:

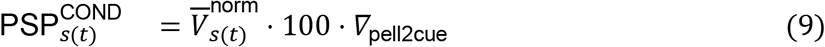

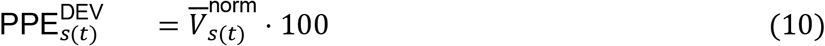

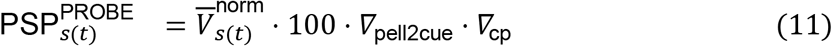

*∇*_pell2cue_ accounts for the difference in the impact of a secondary predictor versus a primary reinforcer, and *∇*_cp_ may account for the forgetting of cue values from COND to the PROBE phase. Both factors are in the range [0,1]. An additional factor for the calculation of 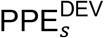 was not necessary. 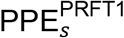 and 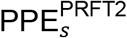 are calculated the same way as 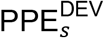.

#### Ha: Outcome-specific encoding deficit

In this version, only the MB system is used, and we assume no forgetting happens from COND to PROBE so *∇*_cp_ is fixed to 1.

We model the inactivation of OFC as implying that the representation of the relevant cues (here, A and B) is potentially only partially distinct. Thus, if, for instance *s*(*t*) = *A* is presented, then writing 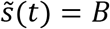 as the ‘other’ cue, we imagine a spillover or fuzziness factor *χ* is introduced that is taken into consideration when doing the updates so that, along with equation 7, we have

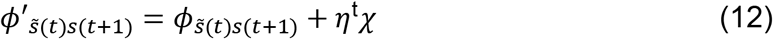

If *χ* = 0, nothing is learned for the opposite state, if *χ* = 1, then exactly the same transition information is learned for both states, and if *χ* > 1, then more is learned for the opposite/unseen state. Note that we continue to consider the outcome pellets to be perfectly distinguishable.

The free parameters used for model fitting are: NegRew, *∇*_pell2cue_, *η*^mb^, *η*^t^ *χ*.

#### Model Fitting

Separate sets of parameters were fit for each animal using scipy.optimize.least_squares, optimizing the mean squared error (MSE) between the real behavioral recordings and the model “behavior” outputs based on the current set of parameters. A weighted MSE was used in order to increase the contribution of the PROBE trials as behavioral differences across groups (control/OFC deactivation) were most apparent here, and the number of trials comparably few (there is 8x more condition trials, so PROBE trials have an 8x higher weight). The following bounds for the parameter fitting were defined as follows:

| Param | NegRew | $\nabla_{\text{pell2cue}}$ | $\eta^{\text{mb}}$ | $\eta^t$ | $\chi$ |
| --- | --- | --- | --- | --- | --- |
| Min | -90 | 0 | 0 | 0 | 0 |
| Max | 0 | 1 | 40 | 40 | 1.5 |

Individual parameter estimates for either of the models were then compared across groups using t-tests and Bonferroni-corrected for multiple comparisons.

#### Parameter Recovery

In order to ensure that recovered parameter values are meaningful in case of the model fits, we checked parameter recoverability. Here, we use known parameter values along with realistic noise to generate synthetic data, and then assess if we can recover from these data values of the parameters that are close to the original generating levels. In order to stay close to the real data, we used the parameters recovered for each animal individually to generate one synthetic dataset/behavioral trace per animal. The noise was generated using individual variability estimates of per trial behavioral measures for each experiment stage (COND, DEV, PROBE). This yields 28 pairs (one pair per animal) of real and estimated parameter values for each of the model’s parameters. Good parameter recoverability is when real and estimated parameter values are well correlated.

Recovery of most of the parameters was good (*r*_NegRew_ = 0.9, *r*_*∇*_pell2cue__ = 0.8, *r*_*η*_mb = 0.8, and *r_χ_* = 0.7); only the recovery of the state transition observation rate *η*^t^ was slightly less faithful (*r*_*η*^t^_ = 0.6), and so should be interpreted cautiously.

Repeating the recovery procedure multiple times produced comparable results. We also used a synthetic generative procedure to assess the posterior correlations between recovered parameter values, something that matters for prediction, albeit less for the overall interpretation of the model. We started out with the median parameter values across animals to generate synthetic data, with noise generated based on the variability of behavioral measures per experiment stage, this time on the group level, and recovered those parameter values from these data. We did this 30 times and assessed the correlations between all pairs of inferred parameters. We found that most of the correlations were mild – although the highest correlations between *∇*_pell2cue_ and *η^t^* (*r* = −0.57), were quite substantial. This is not unexpected, as in effect *∇*_pell2cue_ accounts for the difference between the asymptotic performance at the end of conditioning, which is in turn set by the observation rates.

#### Hypothesis Hb. MB deficit

Hb parameterizes a more conventional view of the effect of OFC inactivation, allowing for a combination between MF and MB learning and control, with the possibility that this combination is disturbed by inactivation.

As hypothesis Hb makes use of both model free and model based value systems, it employs two sets of value distributions: 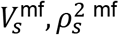 and 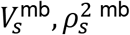. MB learning and inference happens as for hypothesis Ha, except that the imprecision parameter *χ* is not part of Hb. Following Dearden et al. (18), the MF value system uses normal-gamma distributions for characterizing the values 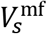 of all states *s*, both terminal (with rewards) and non-terminal (with cues).

For the MF system, each time the animal passes through state s, the value distribution at this state is updated according to either a scalar estimate 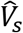 of the long-run reward from that state *s* for the MF system, or the immediate reward *r* using an observation rate *η*^mf^.

Updating the MF values of terminal states is the same as for the MB system (using equation 6) with 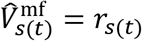 and an observation rate *η* = *η*^mf^. Updating the values of non-terminal (cue) states also follows equation (6), but now (since, in this task, there are no rewards at non-terminal states) with

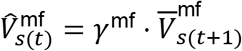

Generally, the estimated value of the model free system is 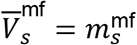 and the estimated variance is given by the expression in equation 3.

According to Hb, both MB and MF contribute to the value of a cue, according to a convex combination parameter *w*^mf^, which is in range [0,1] with 0 meaning only the model-based system is used and 1 that only the model free system is used:

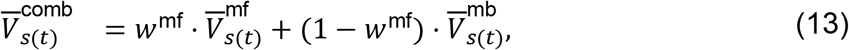

This then generates the normalized value

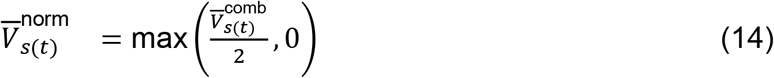

which leads to behavioral measures as in equations (9)-(11).

For convenience of fitting, the observation rate for the transition matrix was fixed to the one for the model based value distributions *η*^t^ = *η*^mb^, and *γ*^mf^ and *γ*^mb^ were set to 1. The free parameters used for model fitting were therefore: NegRew, *∇*_pell2cue_, *∇*_cp_, *η*^mf^, *η*^mb^ and *w*^mf^. As an important simplification, we fixed *w*^mf^ to have the same value for COND and PROBE, even in the inactivation case, as if this had been stamped in during COND, for instance because of heightened MB uncertainty. If *w*^mf^ was lower in PROBE then, we would not have expected such equivalent decreased responding to both cues. An alternative possibility we did not explore is that inactivation would leave the MB system with impaired learning in COND, even at asymptote for both cues; and that if *w*^mf^ was indeed lower in PROBE, reduced responding would come from averaging a persistent value from the MF system with the decreased output of the MB system. This would be an alternative to making parameter *∇*_cp_ small.

The same constraints as above were used for fitting the MB system (albeit with *χ* effectively clamped at 0). Additionally, we had

| Param | $\nabla_{\text{cp}}$ | $\eta^{\text{mf}}$ | $w^{\text{mf}}$ |
| --- | --- | --- | --- |
| Min | 0 | 0 | 0 |
| Max | 1 | 40 | 1 |

Parameter recovery of the observation rate parameters were the least faithful (*r*_*η*^mf^_ = 0.6, *r*_*η*^mb^_ = 0.2 and *r_w_*mf = 06), while the estimated values of the other parameters were closer to real ones (*r*_NegRew_ = 0.9, *r*_**∇**_pell2cue__ = 0.9, *r*_**∇**_cp__ = 0.8). Thus, when interpreting this model, less emphasis should be placed on the first three parameters. Correlations in the recovered values of the parameters were mild – with the highest correlation being between **∇**_pell2cue_ and *w*^mf^ (*r* = −0.54).

## Acknowledgments

We thank Jordi Bonaventura for guidance on chemogenetic methods, Nishika Raheja and Savana Agyemang for technical assistance, as well as Evan Hart and Marios Panayi for stimulating discussions. The opinions expressed in this article are the authors’ own and do not reflect the view of the NIH/DHHS.

## Funding

Intramural Research Program at the National Institute on Drug Abuse (KMC, MG, GS)

Max Planck Society (RS, KL, PD)

Humboldt Foundation (PD).

German Research Foundation grant MA 8509/1-1 (KMC)

## Author contributions

Conceptualization: KMC, MPHG, PD, GS

Methodology: KMC, RS, KL, MPHG, PD, GS

Investigation: KMC

Software: RS, KL, PD

Validation: KMC, RS, KL, PD

Data Curation: KMC, RS

Formal analysis: KMC, RS

Visualization: KMC, RS

Resources: PD, GS

Supervision: PD, GS

Project Management: PD, GS

Writing – original draft: KMC

Writing – review & editing: KMC, RS, PD, GS

## Data and materials availability

all code and data used in this study are available on https://colab.research.google.com/drive/1VYRAnvAO8OmzQpVaJe5radKIZnpEn638?usp=sharing. Additional information on materials and protocols are available upon request to the corresponding authors.

## Supplementary Information

**Table S1.** Statistical results of behavioral experiments.

| **Conditioning (3-way ANOVA)** | **SS** | **MS** | **F (DFn, DFd)** | **P value** |
| --- | --- | --- | --- | --- |
| Sessions | 39952 | 5707 | F (7, 182) = 26.74 | P<0.0001**** |
| (CTRL vs hM4d) | 2512 | 2512 | F (1, 26) = 0.7170 | P=0.4048 |
| (A vs B) | 12.16 | 12.16 | F (1, 26) = 0.02112 | P=0.8856 |
| Sessions x (CTRL vs hM4d) | 6981 | 997.3 | F (7, 182) = 4.672 | P<0.0001**** |
| Sessions x (A vs B) | 874.0 | 124.9 | F (7, 182) = 1.353 | P=0.2279 |
| (CTRL vs hM4d) x (A vs B) | 9.810 | 9.810 | F (1, 26) = 0.01703 | P=0.8972 |
| Sessions x (CTRL vs hM4d) x (A vs B) | 517.3 | 73.90 | F (7, 182) = 0.8008 | P=0.5876 |
| **CTA (3-way ANOVA)** | **SS** | **MS** | **F (DFn, DFd)** | **P value** |
| Sessions | 37224 | 18612 | F (2, 52) = 64.18 | P<0.0001**** |
| (CTRL vs hM4d) | 1344 | 1344 | F (1, 26) = 2.912 | P=0.0998 |
| (ND vs D) | 53832 | 53832 | F (1, 26) = 127.1 | P<0.0001**** |
| Sessions x (CTRL vs hM4d) | 39.38 | 19.69 | F (2, 52) = 0.06789 | P=0.9344 |
| Sessions x (ND vs D) | 41521 | 20761 | F (2, 52) = 83.36 | P<0.0001**** |
| (CTRL vs hM4d) x (ND vs D) | 155.6 | 155.6 | F (1, 26) = 0.3675 | P=0.5497 |
| Sessions x (CTRL vs hM4d) x (ND vs D) | 32.92 | 16.46 | F (2, 52) = 0.06609 | P=0.9361 |
| **PROBE (2-way ANOVA)** | **SS** | **MS** | **F (DFn, DFd)** | **P value** |
| (CTRL vs hM4d) | 806.6 | 806.6 | F (1, 26) = 4.340 | P=0.0472* |
| (A vs B) | 143.3 | 143.3 | F (1, 26) = 1.743 | P=0.1983 |
| (CTRL vs hM4d) x (A vs B) | 658.8 | 658.8 | F (1, 26) = 8.013 | P=0.0088** |

**Table S2.** Comparisons of fitted parameters for the control and hM4d groups withing the two tested reinforcement learning models. Data are represented as mean ± SEM.

| *Ha: Precision deficit model parameters* | **Control** | **hM4d** | **P value** |
| --- | --- | --- | --- |
| *Imprecision term - $\chi$* | 0.64 ± 0.138 | 1.031 ± 0.099 | P=0.027* |
| *Model-based value observation rate - $\eta^{mb}$* | 2.669 ± 1.964 | 5.062 ± 2.626 | P=0.169 |
| *Model-based transition observation rate - $\eta^{tm}$* | 11 ± 4.657 | 22.41 ± 4.477 | P=0.067 |
| *Strength of devaluation - NegRew* | -50.82 ± 5.092 | -60.21 ± 4.81 | P=0.185 |
| *Value adjustment - $\nabla_{pell2cue}$* | 0.718 ± 0.069 | 0.512 ± 0.066 | P=0.04* |

| *Hb: MB vs MF model parameters* | **Control** | **hM4d** | **P value** |
| --- | --- | --- | --- |
| *Model-free observation rate - $\eta^{mf}$* | 3.71 ± 2.778 | 15.21 ± 4.758 | P=0.007** |
| *Model-based observation rate - $\eta^{mb}$* | 12.8 ± 4.51 | 15.77 ± 4.82 | P=0.575 |
| *Contribution of model-free system - $w^{mf}$* | 0.553 ± 0.106 | 0.703 ± 0.081 | P=0.513 |
| *Strength of devaluation - NegRew* | -52.09 ± 5.85 | -54.78 ± 5.243 | P=0.821 |
| *Value adjustment - $\nabla_{pell2cue}$* | 0.761 ± 0.069 | 0.501 ± 0.059 | P=0.012* |
| *Conditioning to probe forgetting - $\nabla_{cp}$* | 0.603 ± 0.079 | 0.385 ± 0.083 | P=0.071 |

**Figure S1.**
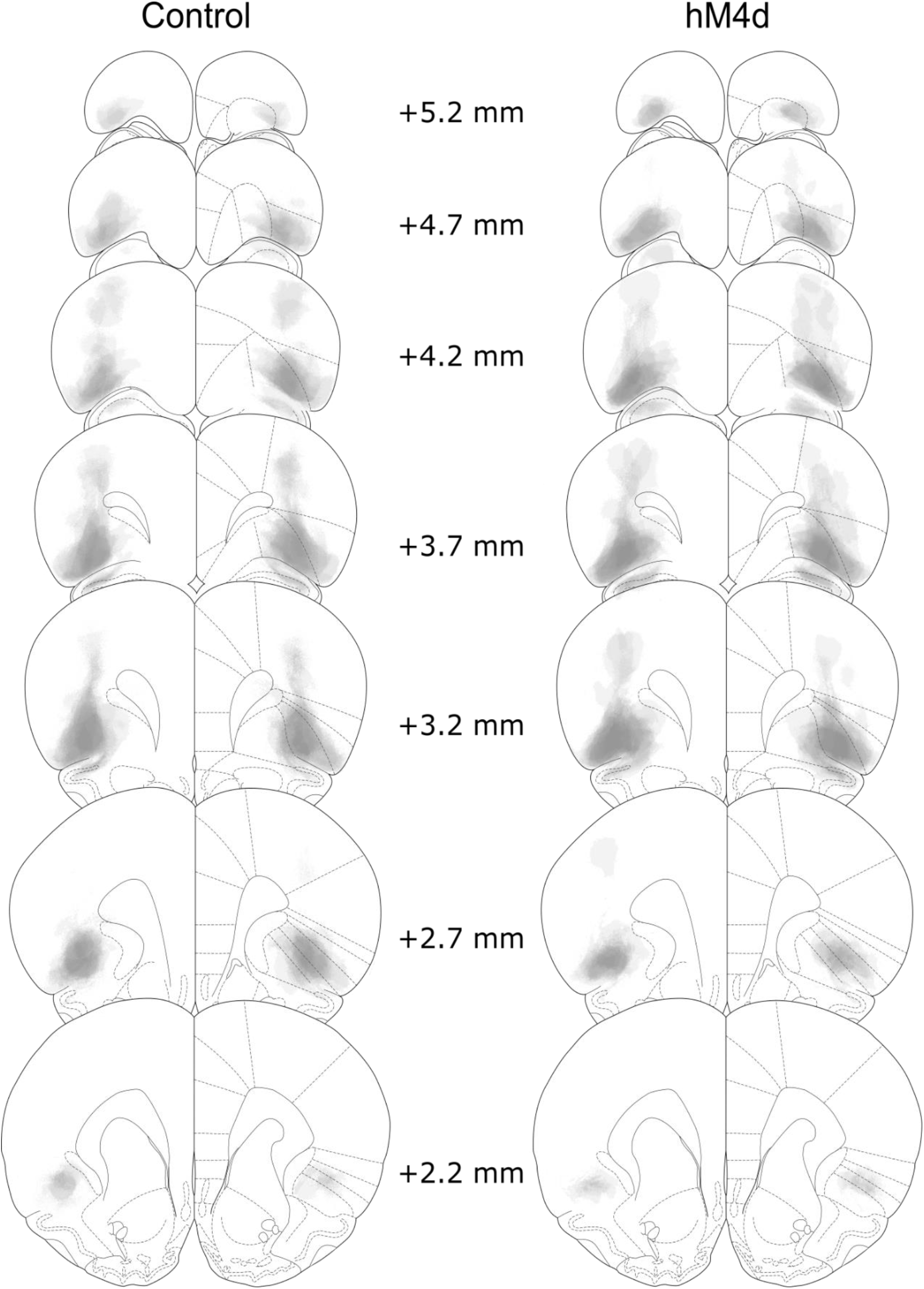
Histological validation of DREADD strategy. Reconstruction of viral expression patterns in the OFC across the control and hM4d groups. Viral spread was mostly contained withing OFC and was similar for control and hM4d subjects.

**Figure S2.**
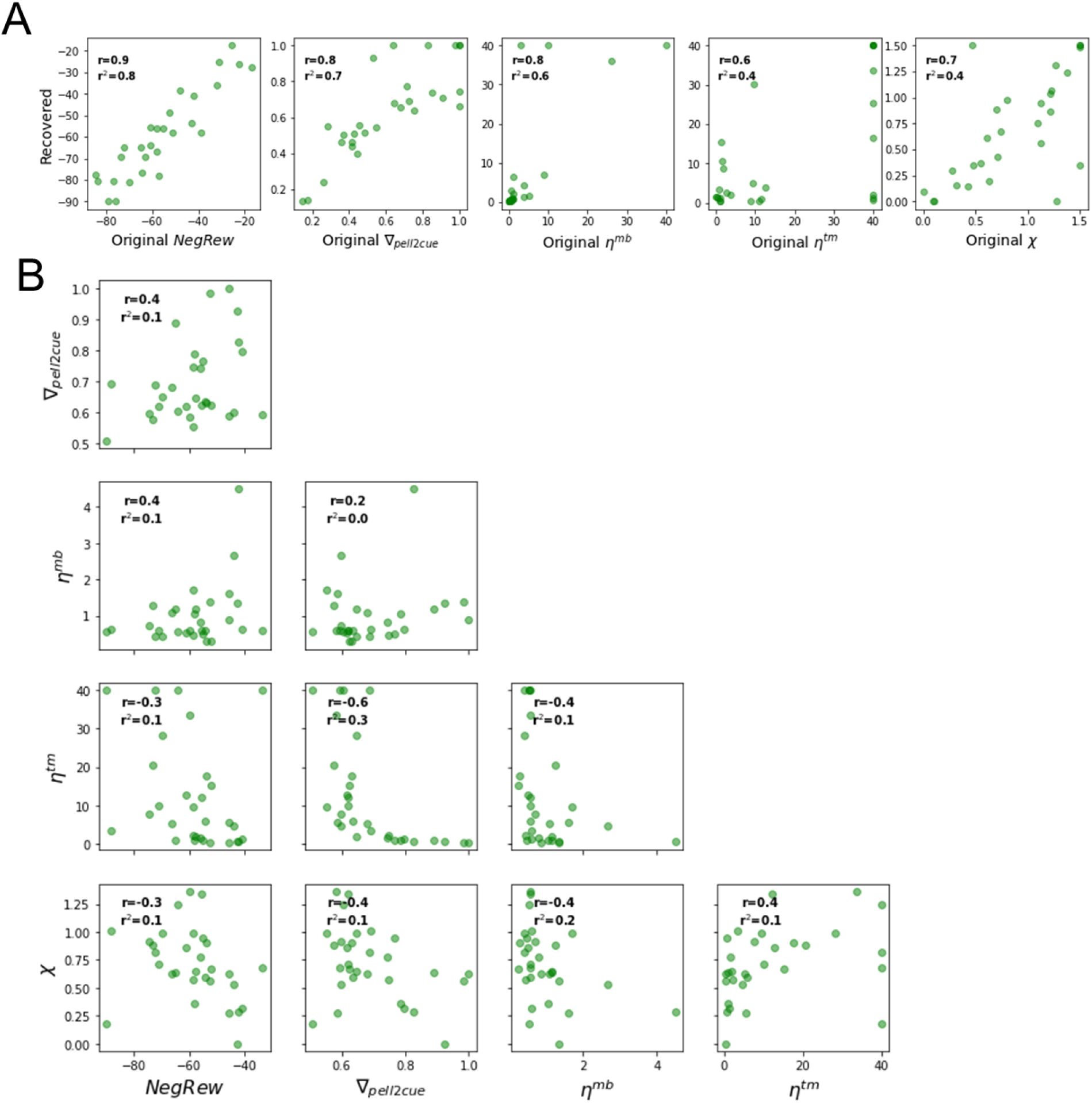
Parameter recovery and correlations for the reinforcement learning model with association specificity deficit. **A:** Correlations between estimated and original parameters. Note that most parameters were recovered with r>0.7, with the least faithfully recovered parameter being the state transition observation rate η^tm^ with r < 0.6. **B:** Correlations between fitted parameters. Note that only correlations between *∇*_pell2cue_ and w^mf^ (*r* = −0.54) in HB and between *∇*_pell2cue_ and *η^tm^* (*r* = −0.57) are substantial.

**Figure S3.**
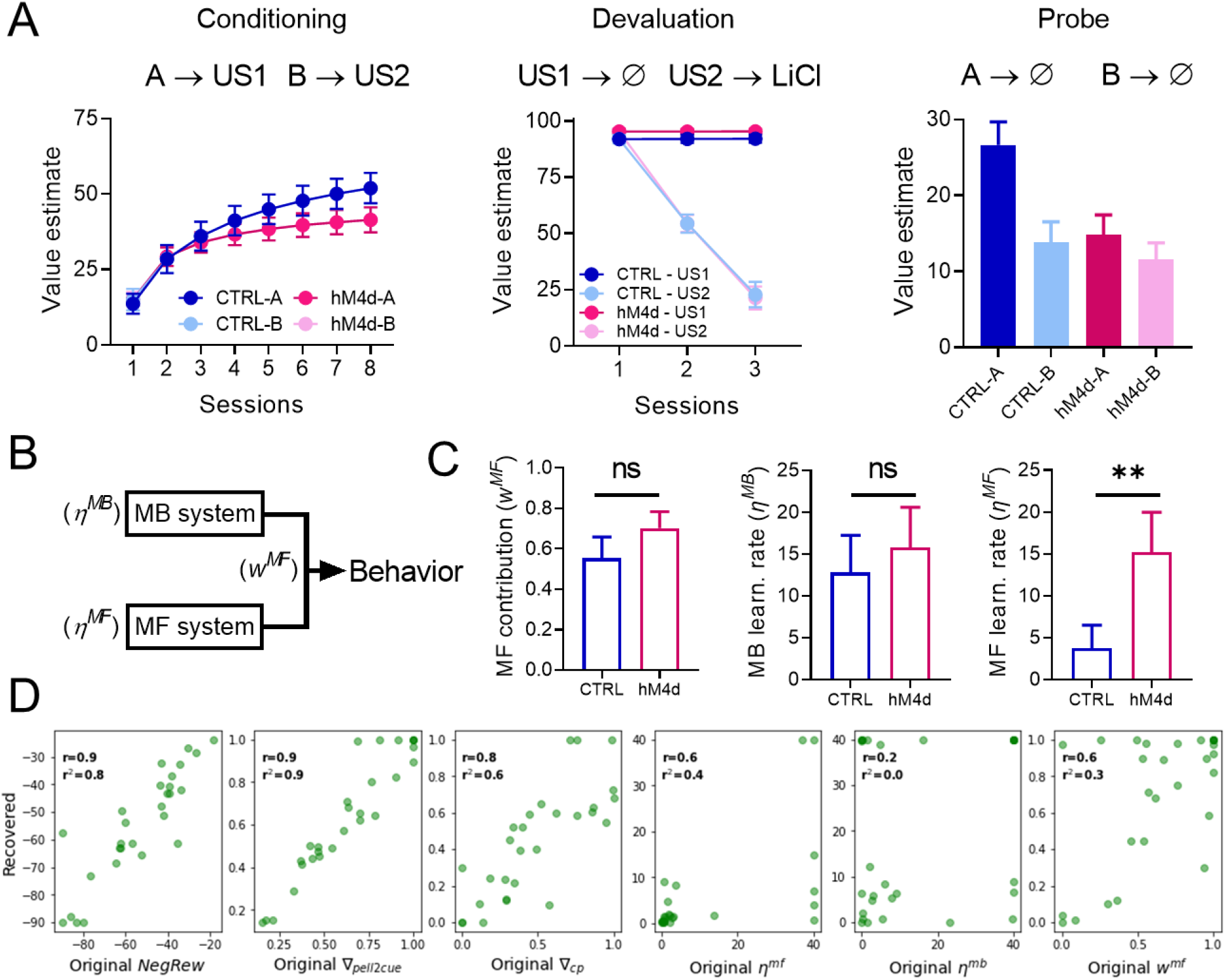
Data fitting with a reinforcement learning model that allows for a shift between model-based (MB) and model-free (MF) learning. (A) Model fit results for our MB vs MF reinforcement learning model. Note that it can also replicate our behavioral results well. (B) Schematic of the critical aspect of the model and the expected result: the observation rate for both the MB and MF systems, as well as the potential contribution of each to behavior, were free parameters, and we expected that the contribution of the MB system would be diminished, either by a reduced MB observation rate or an increase in the MF contribution. (C) Values of the critical observation rate-related parameters, namely the proportion of contribution of the MF (*w*^mf^) system, the MF observation rate (*η*^mf^), and the MB observation rate (*η*^mb^) for both control and hM4d model fits. Note that instead of a reduction in MB learning or proportional contribution, only the MF observation rate was significantly higher in the hM4d group. See table S2 for detailed parameter comparisons. (D) Correlations between estimated and original parameters for the MB vs MF model. Note that parameter recovery of all critical observation rate-related parameters was not very faithful (r < 0.7). Data are represented as mean ± SEM. \*\**P*<0.01.

